# Lipopolysaccharide composition determines the preferred route and entry kinetics of bacterial outer membrane vesicles into host cells

**DOI:** 10.1101/084848

**Authors:** Eloise J O’Donoghue, Douglas F. Browning, Ewa Bielska, Luke Alderwick, Sara Jabbari, Anne Marie Krachler

## Abstract

Outer membrane vesicles are microvesicles shed by Gram-negative bacteria and play important roles in immune priming and disease pathogenesis. However, our current mechanistic understanding of vesicle - host cell interactions is limited by a lack of methods to study the kinetics of vesicle entry and cargo delivery to host cells in real-time. Here, we describe a highly sensitive method to study the kinetics of vesicle entry into host cells in real-time using a genetically encoded probe targeted to vesicles. We found that route of vesicular uptake, and thus entry kinetics and efficiency of cargo release, are determined by the chemical composition of the bacterial lipopolysaccharide. The presence of O-antigen facilitates receptor-independent entry, which enhances both rate and efficiency of cargo uptake by host cells. Collectively, our findings highlight the chemical composition of the bacterial cell wall as a major determinant of secretion-independent delivery of virulence factors during Gram-negative infections.

## INTRODUCTION

Outer membrane vesicles (OMVs) are nano-sized proteoliposomes released from the cell envelope of all Gram negative species of bacteria (Haurat et al, 2015). OMV release is a highly conserved process, occurring under all growth phases and environmental conditions (Bonnington and Kuehn, 2014). OMVs are able to contain and deliver a range of substrates, from large hydrophobic molecules to DNA, making them a versatile and generalised form of secretion that enhances bacterial fitness in hostile environments (Renelli et al, 2004; Kaparakis et al, 2010; Manning and Kuehn, 2011; MacDonald and Kuehn, 2012). They also contribute significantly to pathogenesis, via the delivery of virulence factors such as toxins, adhesins and immunomodulatory compounds directly into the host cell (Alaniz et al, 2007; Lindmark et al, 2009; Roy et al, 2011). In a mouse model, purified OMVs from *Escherichia coli* were sufficient to cause lethal sepsis in the absence of intact bacterial cells, indicating their potency in enhancing infection and inflammatory processes (Park et al, 2010). The immunogenicity and ubiquitous production of OMVs has also led to their successful use in vaccine preparations, such as MeNZB against *Neisseria meningitidis*, representing an application for OMVs in generating immunity against bacterial infections without the risks associated with live cell vaccines (Oster et al, 2007). Whilst many virulence factors have been identified as OMV cargo, the processes underlying their delivery to host cells during infection are not well characterised. Understanding these mechanisms could help to identify targets for inhibition of OMV-associated toxin delivery and lead to attenuation of bacterial infections, as well as helping to achieve their therapeutic potential in medicine, via vaccines and engineered delivery vehicles (Kim et al, 2008; Gujrati et al, 2014; O’Donoghue and Krachler, 2016).

Release of OMVs occurs during infection, and has advantages over other secretion systems. They carry a broad range of cargo, from protein toxins such as VacA and shiga toxin, to hydrophobic molecules such as *Pseudomonas* quinolone signal (PQS), the quorum sensing molecule of *Pseudomonas aeruginosa,* and this cargo is protected from the potentially hostile extracellular environment (Kulp and Kuehn, 2010; Berleman and Auer, 2013). OMV-mediated delivery of virulence factors occurs without requiring close proximity between the bacterial cell and the host cell (Bomberger et al, 2009). The small size of OMVs (20-200 nm) has made studying their interactions with host cells in real time difficult. Previous work has often relied on OMVs labelled with dyes such as fluorescein isothiocyanate (FITC) or dioctadecyloxacarbocyanine perchlorate (DiO). While such dyes allow real time study of OMV entry and cargo delivery processes, the use of membrane labelling of the vesicles may interfere with their physiological characteristics, and alter the mechanism of OMV entry and cargo release (Bauman and Kuehn, 2009; Lulevich et al, 2009; Parker et al, 2010). Other approaches have used immunolabelling of OMV-associated epitopes, such as hemolysin (HlyA) in enterohemorrhagic *Escherichia coli* (EHEC), but this requires fixation of cells at pre-determined time points, and requires assumptions about OMV cargo, which may ignore natural sub-populations of OMVs (Bielaszewska et al, 2013). Some experiments have used host cell phenotypes as an indicator of OMV uptake, such as downstream activation of NF-κB responses in epithelial cells after incubation with OMVs from *Helicobacter pylori* (Kaparakis et al, 2010). However, host cell responses to OMV cargo are decoupled from vesicle uptake and may occur much later than the initial entry event (Wai et al, 2003). The discrepancies in previous observations of OMV entry and cargo delivery may be attributable to the different methods used in these studies, demonstrating the need for an assay that can detect OMV entry processes in a consistent and repeatable manner. Here we describe a novel assay to continuously measure OMV uptake and cargo delivery to host cells with high sensitivity, and in a format that is adaptable for high throughput screening. Using this assay to study entry of EHEC vesicles into host cells, we identified key bacterial and host factors that determine the route of entry, and thereby kinetics and efficiency of vesicular cargo delivery to host cells.

## RESULTS

### A highly sensitive and dynamic assay for monitoring entry of OMV into host cells

We set out to develop an assay that would allow us to monitor entry of OMVs into host cells in real-time. This assay relies on the use of a genetically encoded hybrid reporter probe that is incorporated into the bacterial outer membrane and subsequently targeted to OMVs. ClyA, a cytolysin that is sorted into OMVs by EHEC, acts as the targeting component, and is fused to the catalytic domain of ß-lactamase (Bla), which acts as an enzymatically active probe (Figure 1A). Host cells are incubated with CCF2-AM, a dye composed of a covalently linked coumarin and fluorescein molecule, resulting in FRET and green fluorescence emission specifically in the eukaryotic cytoplasm where it is processed by esterases, while remaining non-fluorescent in its extracellular form. Cytoplasmic processing also prevents the derivative from exiting the cell, thus trapping the fluorescent derivative in the host cell cytoplasm. When OMVs isolated from the producing bacterial strain enter host cells, their ClyA-Bla fusion cargo is able to cleave CCF2-AM, which abolishes FRET and results in a detectable shift in emission from green (530nm) to blue (460nm) fluorescence (Figure 1A). This shift in emission can be monitored in real time to determine the rate of uptake and delivery of OMV cargo to host cells, by comparing the ratio of blue to green fluorescence intensity as an indicator of OMV-host cell interactions.

**Figure 1.**
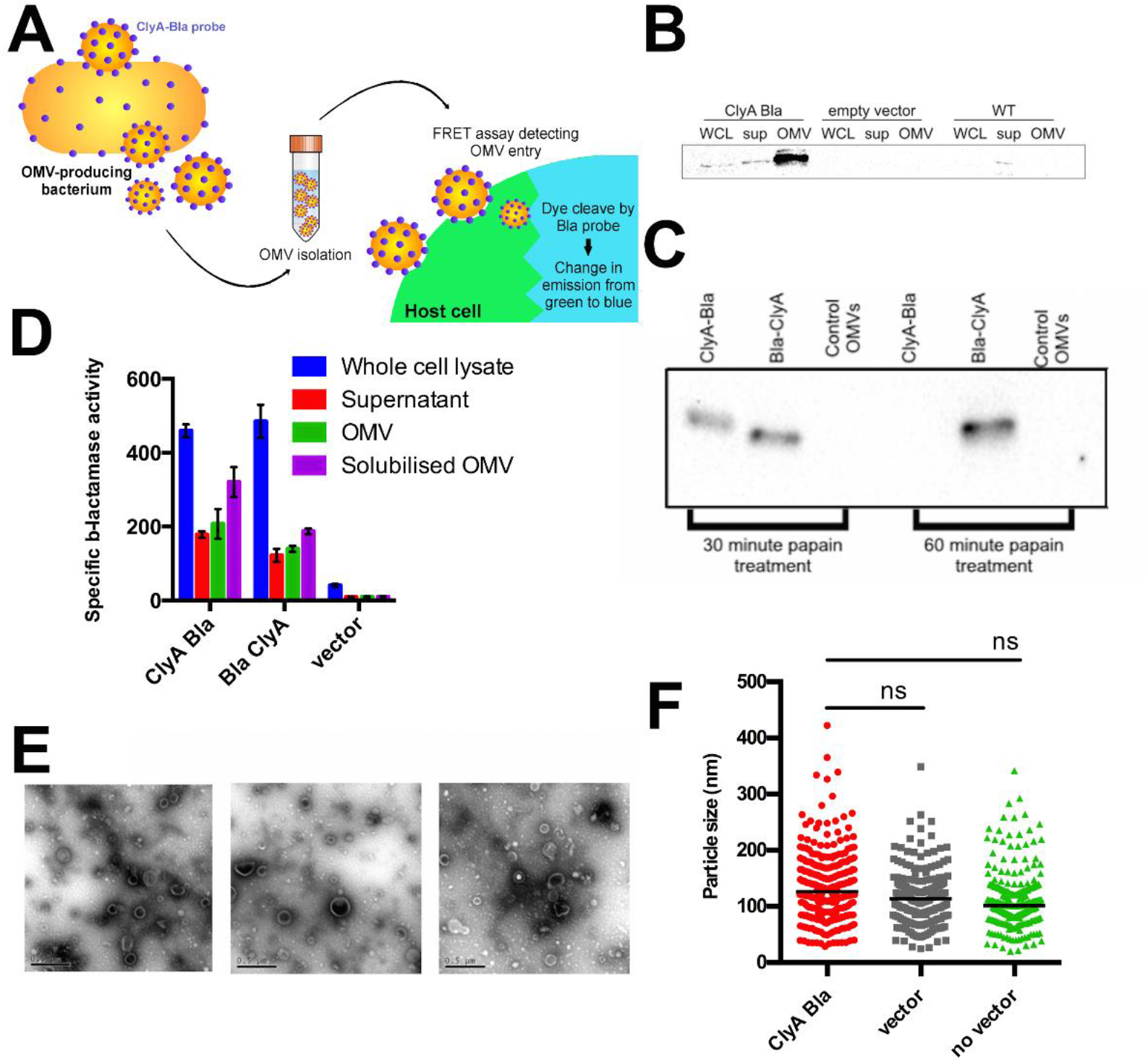
ClyA-Bla probes are targeted to EHEC OMVs and retain their enzymatic activity. (A) Expression of genetically encoded ClyA-Bla probes is induced in bacteria and secreted vesicles are isolated for subsequent experiments. Entry of enzymatically active probes into host cells can be detected using a continuous FRET assay. (B) Whole cell lysate (WCL), supernatant (sup) and outer membrane vesicles (OMV) fractions isolated from EHEC expressing ClyA-Bla, carrying empty vector, or no vector were separated by SDS-PAGE and expression of ClyA-Bla was detected by Western Blotting with α-Bla antibody. (C) OMV fractions from EHEC expressing Cly-Bla, Bla-ClyA or carrying empty vector were treated with papain for 30 or 60 minutes, and used for Western Blotting with α-Bla antibody. (D) Specific enzyme activity in whole cell lysate, supernatant, OMV or solubilized OMV fractions isolated from EHEC expressing ClyA-Bla, Bla-ClyA, or carrying empty vector. (E) Electron micrographs of negative stained OMV fractions from EHEC wt or EHEC ClyA-Bla. Scale bars, 0.5 μm. (F) Nanoparticle tracking analysis of OMV fractions from EHEC expressing ClyA-Bla (red), carrying empty vector (grey) or no vector (green). Results are means ± stdev from at least 200 tracks per sample. Significance was determined using analysis of variance (ANOVA), with a Brown Forsythe test to determine equal variance. NS indicates no significant differences between samples (p value > 0.05).

### ClyA-Bla probes are targeted to EHEC OMVs and retain their enzymatic activity

First, we set out to verify whether ClyA-Bla fusion constructs retained the ability of ClyA to partition into vesicles, and were indeed targeted to EHEC OMVs. Following induction of ClyA-Bla fusion protein production, OMVs were isolated from EHEC expressing either ClyA-Bla (C-terminal fusion) or Bla-ClyA (N-terminal fusion). Western blotting and probing of membranes with Bla-specific antibody revealed the presence of a specific band at 69 kDa, corresponding to intact ClyA-Bla fusion protein in samples from EHEC whole cell lysate, supernatant and OMVs following induction, suggesting that the fusion protein was targeted to and enriched in OMV fractions, as previously reported for non-pathogenic *E. coli* (Kim et al, 2008), (Figure 1B). Protease protection experiments revealed that the ClyA-Bla probe was oriented with Bla facing the exterior of the OMV, as β-lactamase detection by Western blotting was abolished after prolonged treatment of ClyA-Bla OMVs with papain, while the signal remained stable in OMVs containing Bla-ClyA where Bla faces the vesicle lumen (Figure 1C). Enzymatic activity of the probes was assessed by their ability to hydrolyse the β-lactamase substrate nitrocefin. The specific enzyme activity was high for ClyA-Bla OMVs and lower for Bla-ClyA, and both activities were equalized by lysis of vesicles and probe solubilization, suggesting efficient expression of active β-lactamase with the anticipated orientation (inward facing for Bla-ClyA, outward facing for ClyA-Bla) in the isolated OMV fractions (Figure 1D).

Transmission electron microscopy and nanoparticle tracking analysis confirmed that incorporation of ClyA-Bla probes into OMVs had no observable effect on OMV morphology or size (Figure 1E, F and S1). Average OMV concentration was 5 x 10^12^ particles per ml, with sizes ranging from 10-400 nm in diameter, with a mean diameter of 134 nm. The size ranges were in accordance with data obtained previously for OMVs from *E. coli* (Kim et al, 2008) and particle concentrations of all samples were normalized to give a consistent OMV concentration for subsequent experiments.

### OMV-targeted enzyme probes report on vesicle uptake and dismantling by host cells

Having verified the correct targeting, orientation and enzymatic activity of the ClyA-Bla probes, we set out to use them to dissect the uptake and release of vesicle luminal contents in epithelial cells. Due to their different probe orientation, ClyA-Bla OMVs report on uptake of OMVs into host cells, while Bla-ClyA OMVs report on exposure of luminal cargo, or dismantling of OMVs within host cells. Hela epithelial cells loaded with CCF2-AM and exposed to EHEC ClyA-Bla OMVs at a multiplicity of infection (MOI) of 1000 showed a rapid increase in FRET (and thus OMV uptake) over the course of a 3 hour experiment, while Bla-ClyA caused a slower increase in FRET with a notably later onset compared to ClyA-Bla. OMVs devoid of probe did not cause a significant change in FRET (Figure 2A). Despite the different rates in FRET change, the total decrease in FRET after 3 hours (i.e., the efficiency of OMV uptake and cargo release, respectively) was similar for ClyA-Bla and Bla-ClyA, indicating both cargo uptake and vesicle dismantling were complete after 3 hours (Figure 2D). The rates of FRET change were further analyzed by fitting to a cubic spline function and estimating gradients. While the rate of cargo release is stable throughout the experiment (approx. 0.002 s^−1^), the rate of uptake is initially high (approx. 0.007 s^−1^) but gradually decreases and approaches the rate of cargo release (Figure 2B and C).

Confocal fluorescence microscopy of HeLa cells infected with EHEC ClyA-Bla OMVs at an MOI of 1000 also showed a distinct shift from green to blue fluorescence over the course of the experiment compared to uninfected cells, validating the spectral traces collected on the plate reader (Figure 2E). Transmission electron microscopy of Hela cells incubated with OMVs revealed the presence of both external and internalized OMVs (Figure 2F).

**Figure 2.**
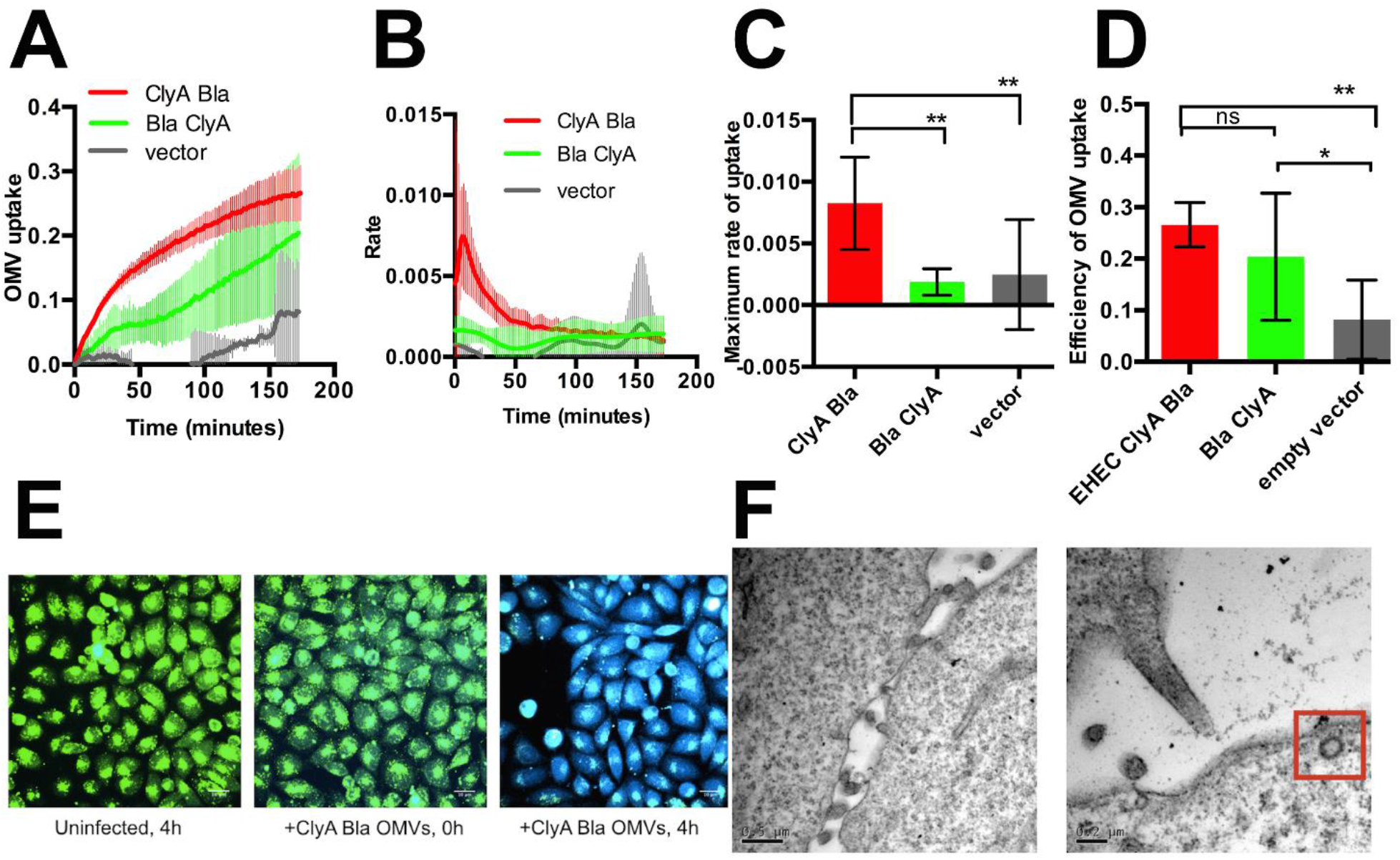
Reporter OMVs enter host cells, and capture dynamics of vesicle uptake and dismantling in real time. (A) CCF2-AM loaded Hela cells were exposed to OMVs from EHEC carrying ClyA-Bla (red), Bla-ClyA (green) or empty vector (grey) at an MOI of 1000 for 3 hours. The FRET signal (ratio of blue:green fluorescence) over time was plotted as mean ± stdev (n=3). (B) polynomials were fitted to each data set using the cubic spline function csaps in Matlab. Numerical estimates of the gradients of the resulting polynomials were determined using the gradient function. Data shown are means ± stdev (n=3). (C) Maximum rates were determined from data in (B) to visualize speed of uptake/cargo release. Data shown are means ± stdev (n=3) and significance was determined by analysis of variance (ANOVA), with a Brown Forsythe test to determine equal variance. (**) indicates p≤0.01. (D) Total FRET changes after 3 hrs were determined from data in (A) and plotted to visualize overall efficiency of uptake/cargo release from OMVs. Data shown are means ± stdev (n=3) and significance was determined by analysis of variance (ANOVA), with a Brown Forsythe test to determine equal variance. (*) indicates p≤0.05, and ns (not significant) p≥0.05. (E) CCF2-AM loaded Hela cells were imaged by confocal microscopy either uninfected (left), following infection with EHEC ClyA-Bla OMVs at an MOI of 1000 immediately (middle) or after 4 hrs of infection (right panel). Images are merged blue/green fluorescence channels and are representative of three independent experiments and 5 fields of view. Scale bar, 10 μm. (F) Following incubation with EHEC ClyA-Bla OMVs for 1 hr, Hela cells were visualized by TEM, and OMVs were observed both externally and internally (red box).

### EHEC OMVs enter host cells more rapidly and efficiently than OMVs from non-pathogenic *E. coli*

Next, we compared the uptake kinetics of OMVs isolated from EHEC and the non-pathogenic *E. coli* strain MG1655. Uptake of EHEC OMVs was faster, compared to MG1655 OMVs (Figure 3A); the maximal rate was higher (Figure 3B-C), and a high rate of uptake was sustained for longer than for the non-pathogenic strain (Figure 3B). The total change in FRET signal was significantly higher, indicating that the overall efficiency of OMV uptake was higher for EHEC than MG1655 (Figure 3D). The higher rate and efficiency of uptake of EHEC OMVs compared to non-pathogenic OMVs were independent of OMV size, which was equivalent, or OMV concentration which was normalized for both strains (Figure S1). Uptake rate and efficiency increased with OMV concentration for both EHEC and MG1655, but the maximal rate plateaued at a lower concentration and lower rate for MG1655 (Figure 3E, F). Taken together, these results suggest EHEC OMVs contain factors absent from MG1655 OMVs that accelerate the rate and increase the efficiency of vesicle uptake by host cells.

**Figure 3.**
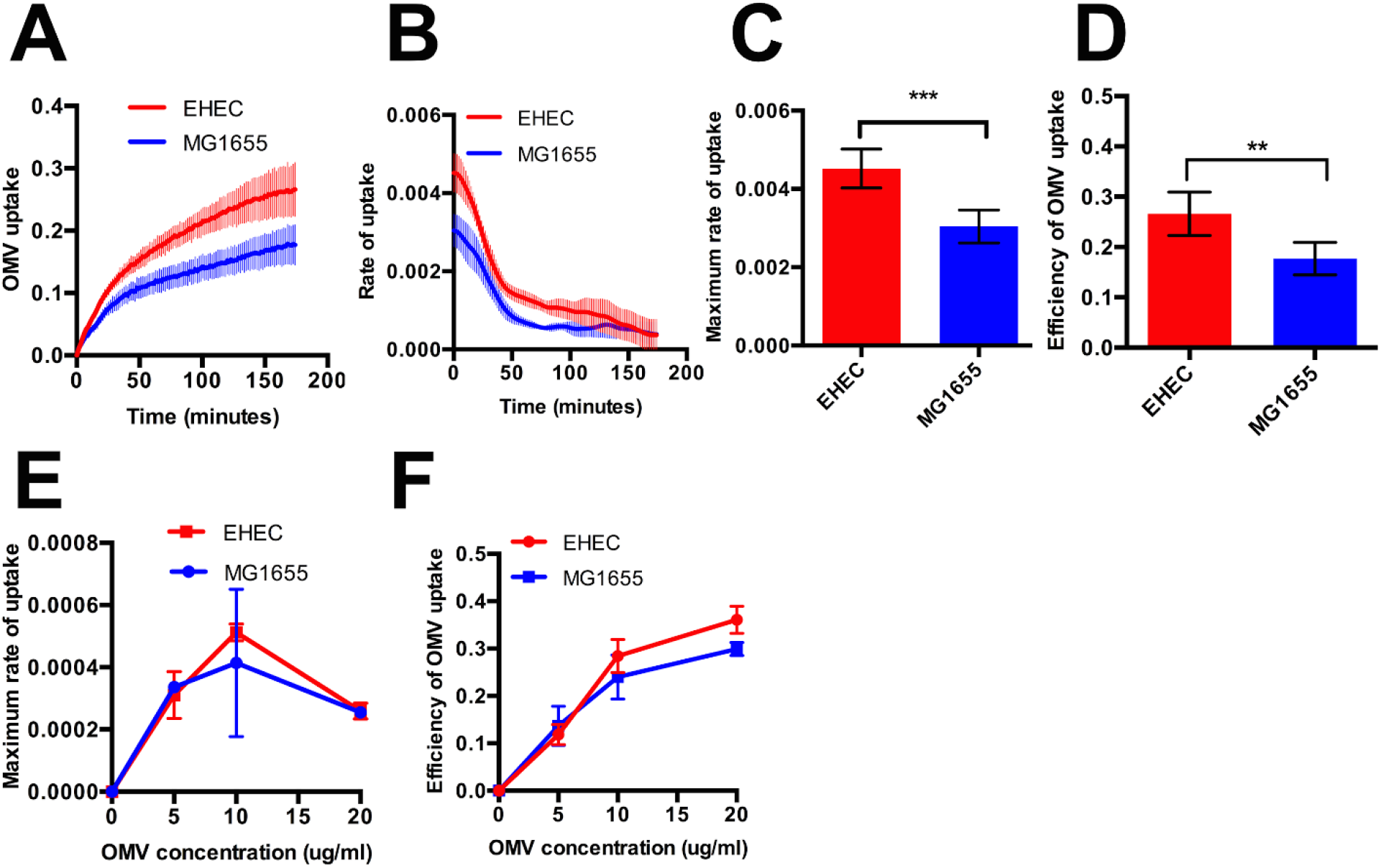
EHEC OMVs enter host cells more rapidly and efficiently than OMVs from E. coli MG1655. (A) CCF2-AM loaded Hela cells were exposed to OMVs from EHEC carrying ClyA-Bla (red), or *E. coli* MG1655 carrying ClyA-Bla (blue) at an MOI of 1000 for 3 hours. The FRET signal (ratio of blue:green fluorescence) over time was plotted as mean ± stdev (n=3). (B) Polynomials were fitted to each data set from (A) using the cubic spline function csaps in Matlab. Numerical estimates of the gradients of the resulting polynomials were determined using the gradient function and plotted as means ± stdev (n=3). (C) Maximum rates were determined from data in (B) to visualize speed of uptake for EHEC (red) or MG1655 (blue) OMVs. Data shown are means ± stdev (n=3) and significance was determined using student’s t-test. (***) indicates p≤0.001. (D) Total FRET changes after 3 hrs were determined from data in (A) and plotted to visualize overall efficiency of uptake for EHEC (red) and MG1655 (blue) OMVs. Data shown are means ± stdev (n=3) and significance was determined by student’s t-test. (**) indicates p≤0.01. (E) Experiments were repeated as for (A), and maximum rates determined as described above. The standard OMV concentration used for experiments (A-D) gives an MOI of 1000 and corresponds to 10 μg/ml of protein. Data are means ± stdev (n=3). (F) Efficiency of uptake (total change in FRET signal over 3 hrs) was determined as described above. Data are means ± stdev (n=3) for each OMV concentration.

### Lipopolysaccharide O-antigen increases efficiency of OMV uptake by host cells

Since OMVs are derived from the outer membrane of Gram-negative bacteria, they contain lipopolysaccharides (LPS), (Cahill et al, 2015). Whilst lipid A and the core oligosaccharide regions are well conserved, many species including EHEC contain a highly variable polysaccharide domain known as the O-antigen (Strauss et al, 2009). The O-antigen constitutes the outermost region of LPS, and due to its length of up to 30 nm, likely the first component presented to host cells upon contact (Strauss et al, 2009). These characteristics led us to hypothesize that the O-antigen moiety of EHEC OMVs may initiate their recognition and uptake by host cells.

In order to investigate this, we compared the uptake kinetics of ClyA-Bla vesicles derived from EHEC O157 wild type cells and the EHEC clinical isolate MA6, which lacks the O157 O-antigen (Rump et al, 2010). While MA6 has an intact *rfb* operon, *gne* was found to have a 1310 bp insertion, preventing synthesis of UPD N-acetyl galactosamine, a substrate required for the first residue of the O-antigen repeating oligosaccharide units (Rush et al, 2010; Rump et al, 2010). Complementation with the *gne* gene from EHEC O157 restored the expression of the O-antigen in EHEC MA6. Despite being isolated from a beef source, MA6 has not been implicated in illness, suggesting a deficiency in its ability to cause infection, despite loss of O-antigen having no effect on growth (Rump et al, 2010). ClyA-Bla reporter OMVs derived from MA6 entered host cells significantly less efficiently than those derived from EHEC wild type cells with intact O-antigen, and overall uptake was decreased by approximately 43% (Figure 4D). The maximum rate of entry of the OMVs from MA6 was not significantly reduced compared with O157 OMVs (Figure 4C).

In addition, enzymatic removal of LPS O-antigen from EHEC O157 wild type OMVs was used to compare the effect of LPS composition on OMVs uptake kinetics. Cellulase, which possesses endo-1,4-beta-D-glucanase activity, was used to hydrolyze the glycosidic bond between α-L-fucose and ß-D-glucose in the repeating unit of the O157 O-antigen (Wang and Reeves, 1998) and efficiently removed the O-antigen from EHEC LPS (Figure S2). Cellulase-treated EHEC reporter OMVs entered host cells significantly slower than those from wild type EHEC, and total uptake efficiency was decreased by approximately 63% (Figure 4C).

*E. coli* MG1655 has lost its ability to produce O-antigen due to a disruption in *wbbL* encoding the rhamnosyltransferase required for O-antigen synthesis (Liu and Reeves, 1994). In the MG1655 L9 strain, wild type *wbbL* has been restored, allowing expression of the strain’s original O16 O-antigen (Browning et al, 2013, Figure S2). We carried out FRET assays with Hela cells exposed to ClyA-Bla reporter OMVs harvested from MG1655 and MG1655 L9 strains, to determine if restoration of its O-antigen would impact OMV uptake kinetics (Figure 4E-H). HeLa cells incubated with reporter OMVs from MG1655 L9 showed a significantly higher total change in FRET signal compared with cells incubated with OMVs from MG1655 (Figure 4A, D). There was no significant difference between the maximum rates of uptake, but the rate of uptake decelerated quicker for the MG1655 than the MG1655 L9 strain (Figure 4B, C), suggesting presence of O16 O-antigen on OMVs causes an increase in efficiency of uptake, but does not affect rate of uptake as much as O157 O-antigen. Taken together, these results suggest that the presence of O-antigen increases the uptake efficiency of OMVs, and depending on the chemical composition of the O-antigen, can enhance the rate of uptake or sustain an elevated uptake rate, compared to LPS lacking O-antigen.

**Figure 4.**
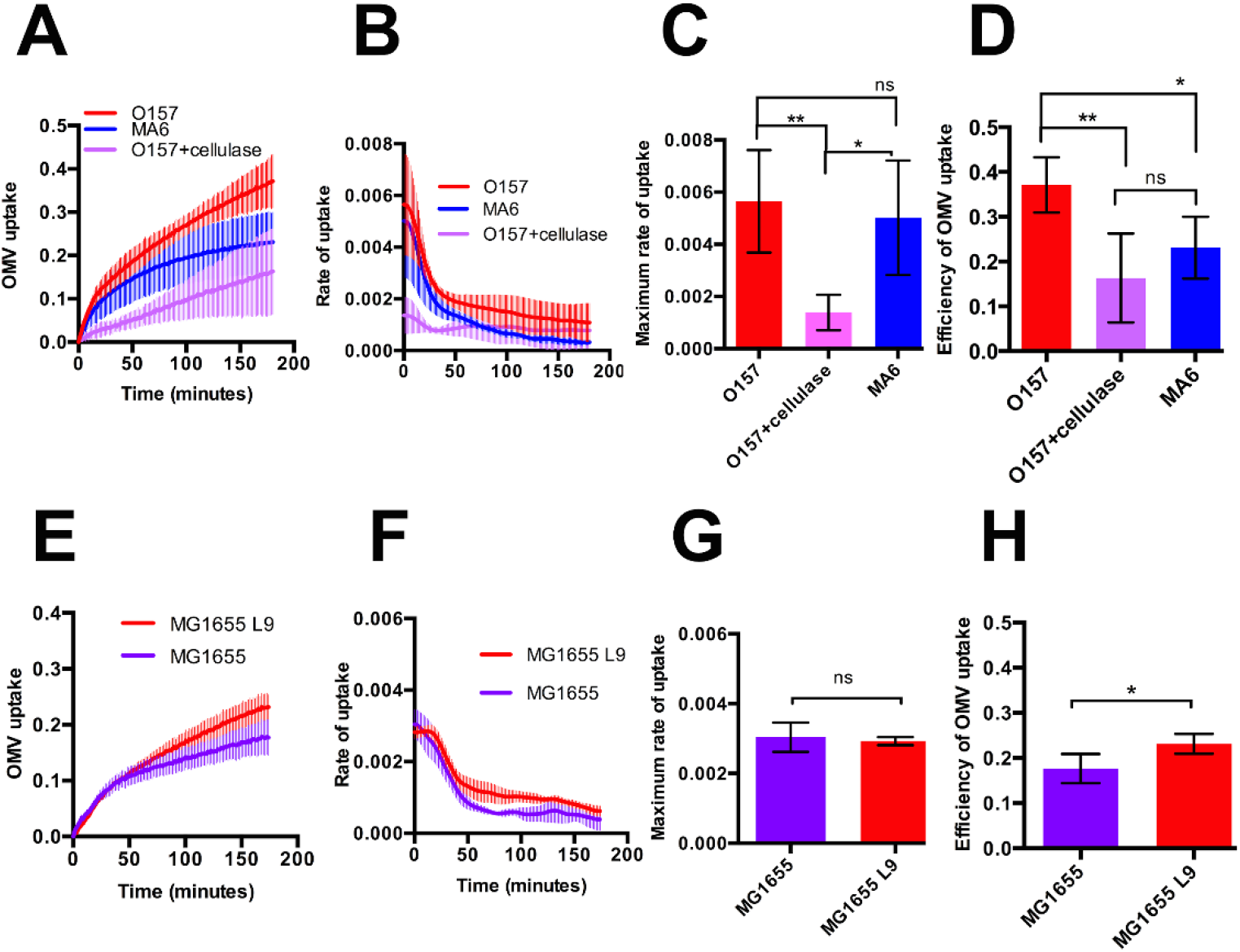
The presence and composition of LPS O-antigen on OMVs affects rate and efficiency of vesicle uptake by host cells. (A) CCF2-AM loaded Hela cells were exposed to ClyA-Bla OMVs isolated from EHEC O157 (red), EHEC MA6 (blue), or EHEC O157 OMVs treated with cellulase (lilac) at an MOI of 1000 for 3 hours. The FRET signal (ratio of blue:green fluorescence) over time was plotted as mean ± stdev (n=3). (B) Polynomials were fitted to each data set from (A) using the cubic spline function csaps in Matlab. Numerical estimates of the gradients of the resulting polynomials were determined using the gradient function and plotted as means ± stdev (n=3). (C) Maximum rates were determined from data in (B) to visualize speed of uptake for EHEC O157 (red), EHEC MA6 (blue) or cellulase-treated EHEC O157 OMVs (lilac). Data shown are means ± stdev (n=3) and significance was determined using ANOVA (***) indicates p≤0.001. (D) Total FRET changes after 3 hrs were determined from data in (A) and plotted to visualize overall efficiency of uptake for EHEC wt (red), EHEC MA6 (blue) or cellulase-treated EHEC wt OMVs (lilac). Data shown are means ± stdev (n=3) and significance was determined by ANOVA (**) indicates p≤0.01, (*) p≤0.05, ns – no significant difference (p≥0.05). (E) CCF2-AM loaded Hela cells were exposed to ClyA-Bla OMVs isolated from MG1655 L9 (red) or MG1655 (purple) at an MOI of 1000 for 3 hours. The FRET signal (ratio of blue:green fluorescence) over time was plotted as mean ± stdev (n=3). (F) Polynomials were fitted to each data set from (E) using the cubic spline function csaps in Matlab. Numerical estimates of the gradients of the resulting polynomials were determined using the gradient function and plotted as means ± stdev (n=3). (G) Maximum rates were determined from data in (F) to visualize speed of uptake for OMVs isolated from MG1655 L9 (red) or MG1655 (purple). Data shown are means ± stdev (n=3) and significance was determined using student’s t-test. ns-no significant differences. (H) Total FRET changes after 3 hrs were determined from data in (E) and plotted to visualize overall efficiency of uptake for OMVs isolated from MG1655 L9 (red) or MG1655 (purple). Data shown are means ± stdev (n=3) and significance was determined by student’s t-test. (*) indicates p≤0.05.

### Macropinocytosis is a minor route of uptake and decreases the efficiency of OMV entry to host cells

Macropinocytosis (Kaparakis-Liaskos and Ferrero, 2015), clathrin dependent endocytosis (Parker et al, 2010), and non-clathrin mediated endocytosis via lipid rafts (Thay et al, 2014; Mondal et al, 2016) have all been proposed as mechanism of vesicle uptake by host cells. These discrepancies may in part be due to differences in OMV composition, as well as methods used to study uptake (O’Donoghue and Krachler, 2016). We used reporter OMVs to evaluate the relative contribution of cellular trafficking pathways to OMV uptake and determine if these are affected by LPS composition. To evaluate the relative contribution of macropinocytosis, OMV uptake kinetics were studied in untreated cells and compared to Hela cells treated with 20 uM blebbistatin to inhibit macropinocytosis (Jiang et al, 2010). Inhibition of macropinocytosis slightly enhanced the rate and overall efficiency of uptake, both for EHEC and MG1655 OMVs (Figure 5), suggesting that a small fraction of OMVs usually enters cells by macropinocytosis but that this pathway is not the major route of uptake, and is a slower way of uptake compared to the major route of entry.

To evaluate whether OMV uptake was dynamin dependent, we compared FRET traces for Hela cells left untreated or treated with 80 uM dynasore to inhibit dynamin-dependent uptake (Nagasawa et al, 2014) prior to incubation with EHEC or MG1655 OMVs. Dynasore treatment almost completely abolished uptake of both EHEC and MG1655 OMVs (Figure 5), suggesting both species use either receptor-mediated endocytosis or caveolin-mediated endocytosis as their major uptake route.

**Figure 5.**
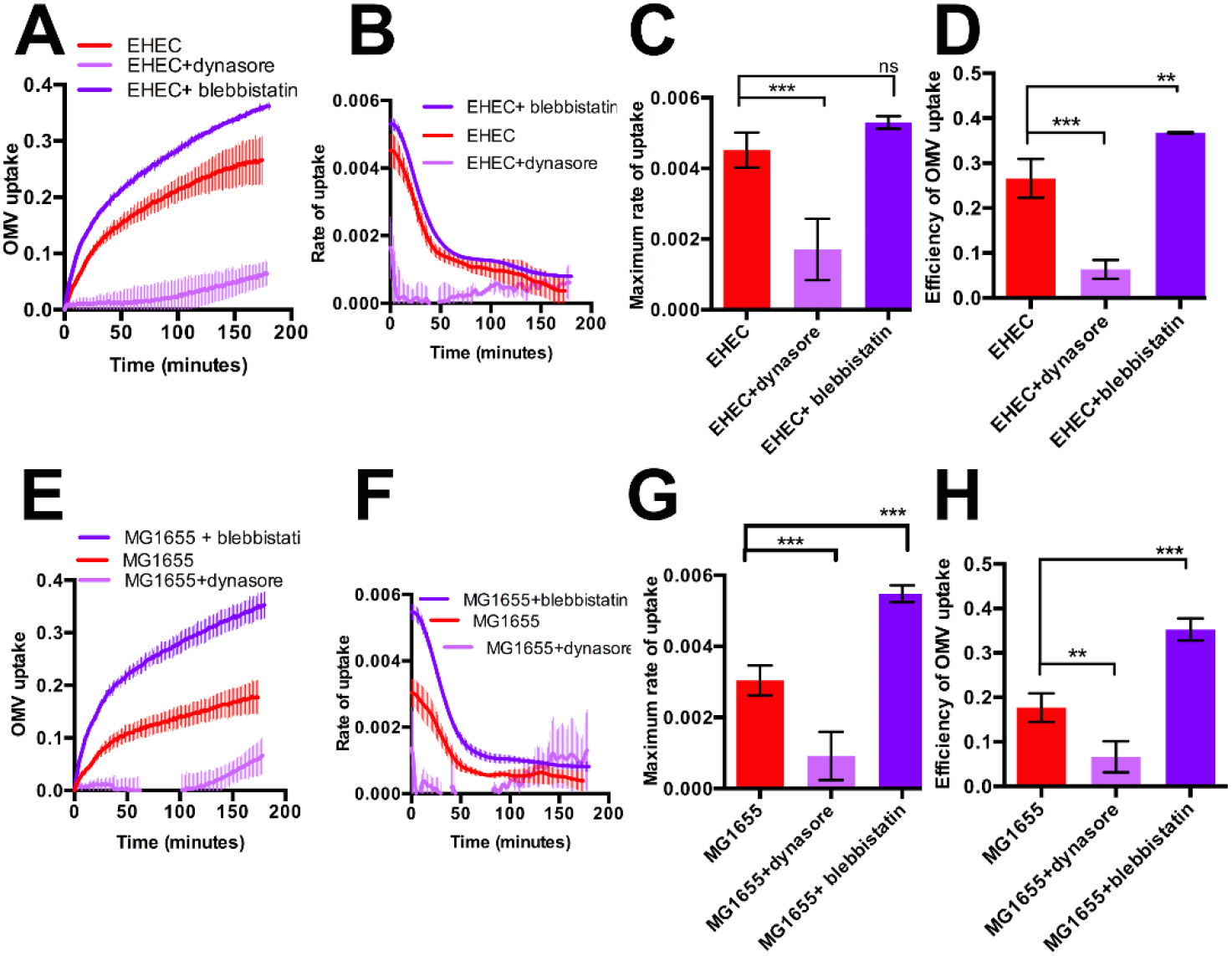
Macropinocytosis is a minor route of uptake and decreases the efficiency of OMV entry to host cells. (A) Hela cells were either left untreated (red), or pre-treated with 80 uM dynasore (lilac) or 20 uM blebbistatin (violet) and exposed to ClyA-Bla OMVs isolated from EHEC at an MOI of 1000 for 3 hours. The FRET signal (ratio of blue:green fluorescence) over time was plotted as mean ± stdev (n=3). (B) Polynomials were fitted to each data set from (A) using the cubic spline function csaps in Matlab. Numerical estimates of the gradients of the resulting polynomials were determined using the gradient function and plotted as means ± stdev (n=3). (C) Maximum rates were determined from data in (B) to visualize speed of OMV uptake into untreated Hela cells (red), or cells pre-treated with 80 uM dynasore (lilac) or 20 uM blebbistatin (violet). Data shown are means ± stdev (n=3) and significance was determined using ANOVA (****) indicates p≤0.0001, ns-not significant (p≥0.05). (D) Total FRET changes after 3 hrs were determined from data in (A) and plotted to visualize overall efficiency of uptake into untreated Hela cells (red), or cells pre-treated with 80 uM dynasore (lilac) or 20 uM blebbistatin (violet). Data shown are means ± stdev (n=3) and significance was determined by ANOVA (***) indicates p≤0.001, (**)p≤0.01. (E) Hela cells were either left untreated (red), or pre-treated with 80 uM dynasore (lilac) or 20 uM blebbistatin (violet) and exposed to ClyA-Bla OMVs isolated from MG1655 at an MOI of 1000 for 3 hours. The FRET signal (ratio of blue:green fluorescence) over time was plotted as mean ± stdev (n=3). (F) Polynomials were fitted to each data set from (E) using the cubic spline function csaps in Matlab. Numerical estimates of the gradients of the resulting polynomials were determined using the gradient function and plotted as means ± stdev (n=3). (G) Maximum rates were determined from data in (F) to visualize speed of MG1655 OMV uptake into untreated Hela cells (red), or cells pre-treated with 80 uM dynasore (lilac) or 20 uM blebbistatin (violet). Data shown are means ± stdev (n=3) and significance was determined using ANOVA (***) indicates p≤0.001. (H) Total FRET changes after 3 hrs were determined from data in (E) and plotted to visualize overall efficiency of MG1655 OMV uptake into untreated Hela cells (red), or cells pre-treated with 80 uM dynasore (lilac) or 20 uM blebbistatin (violet). Data shown are means ± stdev (n=3) and significance was determined by ANOVA (***) indicates p≤0.001, (**) p≤0.01.

### Chemical composition of LPS determines the preferred route of vesicular uptake by host cells

Since our previous experiments had revealed entry of OMVs into host cells was dynamin - dependent, we next sought to determine whether the entry of OMVs was dependent on clathrin-mediated endocytosis, which utilises protein receptors on the host cell surface, or lipid raft-mediated endocytosis, both of which require dynamin (Vercauteren et al, 2010, Chatterjee and Chaudhuri, 2011; Mulcahy et al, 2014). To inhibit clathrin-mediated endocytosis, protein receptors on host cells were removed by treatment with 5 μg/ml papain prior to incubation with EHEC, MG1655 or MG1655 L9 OMVs. Removal of protein receptors from the host surface led to a marked increase in both rate and efficiency of uptake for EHEC and MG1655 L9 OMVs, but abolished uptake of MG1655 OMVs (Figure 6A-L). Pre-treatment of host cells with 1 μg/ml chlorpromazine, which disrupts the formation of clathrin-coated pits (Wang, Rothberg and Anderson, 1993), had a similar effect to papain treatment, and increased rate and efficiency of uptake for EHEC and MG1655 L9, but not for MG1655 OMVs (Figure 6). This indicates that OMVs from MG1655, which lack O-antigen, depend on receptor-mediated endocytosis to enter host cells, whereas EHEC and MG166 L9 OMVs, which feature intact O-antigen, are able to utilize receptor-independent pathways as their major route of entry. While a small fraction of EHEC vesicles appears to enter cells via receptor-mediated endocytosis, inhibition of this route actually increases their uptake rate and efficiency.

### OMVs from EHEC preferentially use lipid raft mediated endocytosis to enter host cells

Since entry of OMVs containing intact O-antigen seemed to be receptor-independent, we investigated whether these OMVs required lipid rafts to enter host cells. Lipid raft-mediated endocytosis was inhibited using 5 mM methyl-ß-cyclodextrin treatment to sequester cholesterol from host cell membranes. Alternatively, cells were treated with 1 μg/ml filipin to disrupt the dynamics of cholesterol-rich microdomains (Nagasawa et al, 2014) prior to incubation with reporter OMVs. Disruption of cholesterol-rich microdomains via both methods led to a marked reduction in rate and efficiency of OMV uptake, and this reduction was most pronounced for EHEC OMVs. Filipin in particular almost completely abolished the uptake of EHEC OMVs (Figure 6A-D). Lipid raft domains can also contain caveolins, which coat cave-like invaginations in the plasma membrane and result in internalization of contents in a process that is also dynamin-dependent (Rewatkar et al, 2015). Since our previous experiments demonstrated the requirement of dynamin for OMV entry (Figure 5), we conclude lipid-raft dependent, caveolin-mediated endocytosis is the major route of entry for EHEC OMVs.

**Figure 6.**
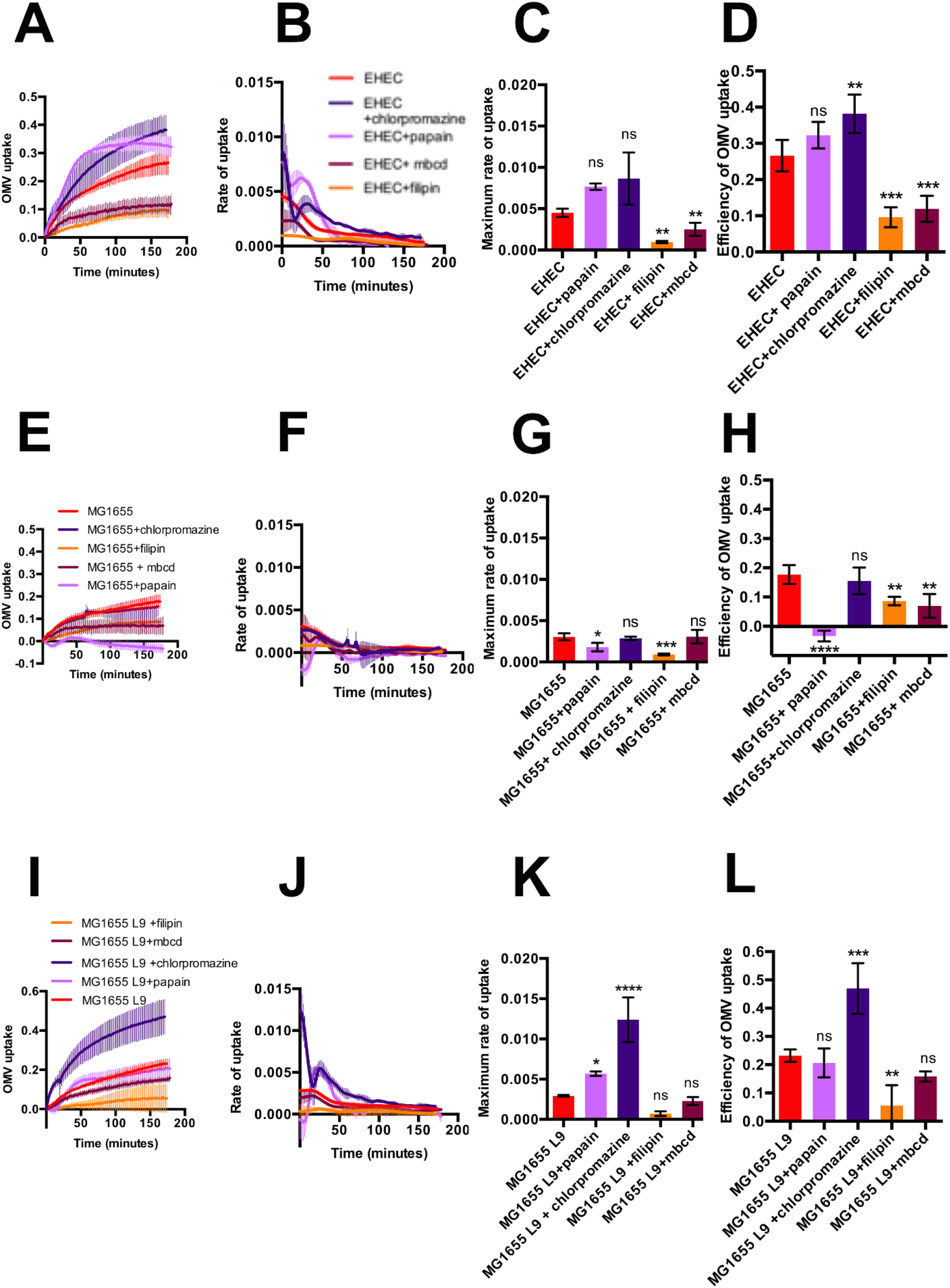
OMVs lacking O-antigen rely on receptor-mediated endocytosis, while OMVs featuring O-antigen preferentially enter host cells via lipid rafts. (A) Hela cells were either left untreated (red), or pre-treated with 5 ug/ml papain (lilac), 1 ug/ml chlorpromazine (purple), 5mM methyl-b cyclodextrin (maroon) or 1μg/ml filipin (orange) and exposed to ClyA-Bla OMVs isolated from EHEC at an MOI of 1000 for 3 hours. The FRET signal (ratio of blue:green fluorescence) over time was plotted as mean ± stdev (n=3). (B) Polynomials were fitted to each data set from (A) using the cubic spline function csaps in Matlab. Numerical estimates of the gradients of the resulting polynomials were determined using the gradient function and plotted as means ± stdev (n=3). (C) Maximum rates were determined from data in (B) to visualize speed of OMV uptake into untreated Hela cells (red), or cells pre-treated with 5 ug/ml papain (lilac), 1 ug/ml chlorpromazine (purple), 5mM methyl-b cyclodextrin (maroon) or 1μg/ml filipin (orange). Data shown are means ± stdev (n=3) and significance was determined using ANOVA (**) indicates p≤0.01, ns-not significant (p≥0.05). (D) Total FRET changes after 3 hrs were determined from data in (A) and plotted to visualize overall efficiency of uptake into untreated Hela cells (red), or cells pre-treated with 5 ug/ml papain (lilac), 1 ug/ml chlorpromazine (purple), 5mM methyl-b cyclodextrin (maroon) or 1μg/ml filipin (orange). Data shown are means ± stdev (n=3) and significance was determined by ANOVA (***) indicates p≤0.001, (**) p≤0.01. (E) Hela cells were either left untreated (red), or pre-treated with 5 ug/ml papain (lilac), 1 ug/ml chlorpromazine (purple), 5mM methyl-b cyclodextrin (maroon) or 1μg/ml filipin (orange) and exposed to ClyA-Bla OMVs isolated from MG1655 at an MOI of 1000 for 3 hours. The FRET signal (ratio of blue:green fluorescence) over time was plotted as mean ± stdev (n=3). (F) Polynomials were fitted to each data set from (E) using the cubic spline function csaps in Matlab. Numerical estimates of the gradients of the resulting polynomials were determined using the gradient function and plotted as means ± stdev (n=3). (G) Maximum rates were determined from data in (F) to visualize speed of MG1655 OMV uptake into untreated Hela cells (red), or cells pre-treated with 5 ug/ml papain (lilac), 1 ug/ml chlorpromazine (purple), 5mM methyl-b cyclodextrin (maroon) or 1μg/ml filipin (orange). Data shown are means ± stdev (n=3) and significance was determined using ANOVA (***) indicates p≤0.001, (*) p≤0.05, ns-not significant (p≥0.05). (H) Total FRET changes after 3 hrs were determined from data in (E) and plotted to visualize overall efficiency of MG1655 OMV uptake into untreated Hela cells (red), or cells pre-treated with 5 ug/ml papain (lilac), 1 ug/ml chlorpromazine (purple), 5mM methyl-b cyclodextrin (maroon) or 1μg/ml filipin (orange). Data shown are means ± stdev (n=3) and significance was determined by ANOVA (****) indicates p≤0.0001, (**) p≤0.01. (I) Hela cells were either left untreated (red), or pre-treated with 5 ug/ml papain (lilac), 1 ug/ml chlorpromazine (purple), 5mM methyl-b cyclodextrin (maroon) or 1μg/ml filipin (orange) and exposed to ClyA-Bla OMVs isolated from MG1655 L9 at an MOI of 1000 for 3 hours. The FRET signal (ratio of blue:green fluorescence) over time was plotted as mean ± stdev (n=3). (J) Polynomials were fitted to each data set from (I) using the cubic spline function csaps in Matlab. Numerical estimates of the gradients of the resulting polynomials were determined using the gradient function and plotted as means ± stdev (n=3). (K) Maximum rates were determined from data in (J) to visualize speed of MG1655 L9 OMV uptake into untreated Hela cells (red), or cells pre-treated with 5 ug/ml papain (lilac), 1 ug/ml chlorpromazine (purple), 5mM methyl-b cyclodextrin (maroon) or 1μg/ml filipin (orange). Data shown are means ± stdev (n=3) and significance was determined using ANOVA (****) indicates p≤0.0001, (*) p≤0.05, ns-not significant (p≥0.05). (L) Total FRET changes after 3 hrs were determined from data in (I) and plotted to visualize overall efficiency of MG1655 L9 OMV uptake into untreated Hela cells (red), or cells pre-treated with 5 ug/ml papain (lilac), 1 ug/ml chlorpromazine (purple), 5mM methyl-b cyclodextrin (maroon) or 1μg/ml filipin (orange). Data shown are means ± stdev (n=3) and significance was determined by ANOVA (***) indicates p≤0.001, (**) p≤0.01.

### Purified LPS competes with EHEC OMVs for caveolin mediated endocytosis

Finally, we considered whether purified LPS would competitively inhibit OMV uptake into host cells. Prior to infection with OMVs, HeLa cells were pre-incubated with 1 μg/ml LPS from either smooth or rough strains (i.e, with or without O-antigen). Supplementation with LPS strongly decreased the rate of uptake of EHEC OMVs, but had no significant effect on uptake of OMVs from MG1655 or L9 (Figure S3). LPS is known to associate with the host TLR4 receptor, but inhibition of TLR4 by pre-treatment of cells with C34 (Neal et al, 2013) did not inhibit uptake of OMVs, suggesting that they are not competing for TLR4 (Figure S4). TLR4 activation also requires CD14 binding, a protein localised in cholesterol rich lipid raft domains (Plociennikowska et al, 2015). We demonstrated that entry of EHEC OMVs is strongly dependent on cholesterol-rich regions of the membrane and we suggest that purified LPS, by recruiting TLR4 signaling complexes in cholesterol-rich regions of the membrane, competes with OMVs for binding in these domains, resulting in a reduction in vesicular uptake.

## DISCUSSION

Interactions between bacterial outer membrane vesicles and epithelial cells are now recognized as an important driver of bacterial pathogenesis, and play key roles in toxin delivery, modulation of mucosal immune responses and immune priming. Yet, our ability to study vesicle-host cell interactions and dissect bacterial and host factors determining route of uptake and thus, the fate of vesicular cargo within the host, has been limited by a lack of methods to study vesicle entry and cargo release in real-time and without altering the physicochemical properties of the vesicle. Here we describe a novel assay that fulfils these requirements and allowed us to study the kinetics of OMV uptake with enough temporal resolution to reveal critical differences between rate and uptake efficiency of vesicles derived from pathogenic and non-pathogenic strains. The assay is sensitive enough to register nuanced differences in the kinetics of OMVs decorated with O-antigens of different chemical composition. Current methods for studying the mechanisms of OMV-mediated delivery of bacterial cargo into host cells have produced inconsistent and often contradictory findings, often relying on less quantitative approaches, such as membrane labelling and immunofluorescence, or using changes in cytotoxicity or host cell phenotype as an indicator of OMV cargo delivery. The method presented here provides a consistent platform to further study the pathways involved and measure the kinetics of OMV entry into host cells by utilising a sensitive, adaptable and quantifiable approach that can easily be adapted to a high-throughput format, and potentially to study OMV-host interactions *in vivo* (Raz et al, 1998).

We selected EHEC as a proof-of-concept species here, since EHEC OMVs have been shown to play a crucial role in toxin stabilization and delivery (Aldick et al, 2008), and have been considered as a means to vaccinate and protect against hemolytic uremic syndrome, a severe complication of EHEC infection (Choi et al, 2014). However, we expect the genetically encoded ClyA-Bla reporter would similarly be targeted to other Gram-negative species of interest, and this will be subject of further investigation to clarify if the herein identified surface features are equally important determinants in driving uptake of OMVs from other species.

Our experiments demonstrated that OMVs from EHEC O157, a successful pathogenic strain of *E. coli*, enter host cells more rapidly and efficiently than those from the lab strain *E. coli* MG1655 (Figure 3). OMVs are able to transport a wide variety of cargo molecules from the bacterial cell into its external environment, and thus the roles that OMVs have in enhancing bacterial fitness are equally diverse (Kuehn and Kesty, 2005; Haurat et al, 2015). This data suggests that pathogens such as EHEC may have fine-tuned their cell wall composition in accordance with the role of OMVs for delivery of virulence factors, such as hemolysins and shiga-like toxins, into the host during infection (Horstmann and Kuehn, 2000; Manning and Kuehn, 2011; Bielaszewska et al, 2013).

Whilst there are many genes that are likely to contribute to the pathogenesis of EHEC, with 1632 proteins present that are absent in MG1655, a significant component of OMVs is LPS, a structure unique to Gram-negative bacteria (Hayashi et al, 2001). Recent work showed that EHEC OMVs allow efficient delivery of LPS into the host cell cytoplasm, resulting in inflammatory responses, caspase-11 activation and cell death (Kunsmann et al, 2015; Vanaja et al, 2016). LPS is composed of the well conserved Lipid A, which forms the outer leaflet of the lipid bilayer and is attached to the core oligosaccharide regions, and in ‘smooth’ strains the core is linked to a highly variable, repetitive polysaccharide region known as O-antigen (Park et al, 2009; Strauss et al, 2009). O-antigen is found at the outer-most portion of the LPS and although variable in length, can consist of up to 100 repeating units and protrude over 30 nm from the membrane (Strauss et al, 2009). The presence of O-antigen can provide several fitness advantages, such as enhancing resistance to complement, phagocytosis and phage infection (Liang-Takasaki et al, 1982; Van der Ley et al, 1986). Pathogenic strains are often ‘smooth’, and the O-antigen can increase bacterial virulence, with the length of the O-antigen positively correlated with the ability of the bacterial cell to adhere to host cells and tissues, while loss of O-antigen results in defects in colonisation, biofilm formation, and increased clearance of infection (Sheng et al, 2008; Strauss et al, 2009; Hathroubi et al, 2016). The importance of O-antigen for adhesion to host epithelial cells has been previously demonstrated in studies using *H. pylori*, but had not been explored in the context of OMVs (Edwards et al, 2000). The position of the O-antigen at the outer surface of the OMV means it may be of relevance for establishing initial contact and facilitating adhesion and entry.

MG1655 is a ‘rough’ strain that has undergone a mutation in the *wbbL* gene of the *rfb* cluster, causing loss of its native O16 O-antigen (Liu and Reeves, 1994). When synthesis of the O16 antigen was restored to generate MG1655 L9, it was able to colonize the intestine of *Caenorhabditis elegans* and cause disease, suggesting that the O-antigen is a requirement for pathogenesis (Browning et al, 2013). In addition, loss of the EHEC O157 antigen results in reduced virulence of the strain in an animal model (Miyashita et al, 2012). In our experiments, OMVs from EHEC MA6, which lacks the O-antigen, entered host cells significantly less efficiently than OMVs from wild type EHEC. This was also observed when the O-antigen subunits were enzymatically cleaved using a glycoside hydrolase. The presence of O16 antigen for MG1655 L9 OMVs did not significantly affect the maximum rate of uptake, but did significantly increase the overall efficiency of uptake compared to MG1655 OMVs by sustaining a higher uptake rate for longer. These differences we observed between O157 and O16 antigens suggest the chemical composition and length of the O-antigen may have a role in fine-tuning entry kinetics and fate of cargo post-uptake. The sugar composition of the O-antigen is highly variable, as is the number of repeating units (Franco, Liu and Reeves, 1998). The O-antigen of most *E. coli* strains has 10-18 repeats, whilst EHEC O157 can exceed 80 repeats (Franco, Liu and Reeves, 1998; Kalynych et al, 2011). The length of the O-antigen in EHEC can be over 30 nm, and the number of repeating units has been previously positively correlated with virulence (Murray, Attridge and Morona, 2006; Strauss et al 2009).

We used our newly-devised assay to identify the relative contributions of cellular uptake pathways to OMV entry into host cells. Inhibition of clathrin-mediated endocytosis by treatment of cells with papain resulted in a significantly increased rate of uptake for EHEC, but a significant reduction in entry for OMVs from MG1655. For OMVs from MG1655 L9, there was no change in rate but an increased total uptake, suggesting a change in the OMV entry phenotype in the presence of O-antigen. This indicates that OMVs without O-antigen rely on protein receptors and clathrin-mediated endocytosis in order to gain access to the host cell, whilst OMVs with O-antigen are not inhibited by the loss of this pathway. We propose that the presence of the O-antigen may lead to possible ligands in the inner regions of the LPS being obscured, whereas these potential receptor binding regions are exposed in the absence of O-antigen, and interestingly, loss of the receptor-mediated pathway is beneficial to EHEC, and this may be due to a preference for the non-receptor mediated pathways which allow for more efficient cargo delivery from OMVs.

When lipid raft mediated endocytosis was inhibited, total uptake of OMVs from all three strains was reduced, but the relative decrease in efficiency was largest for EHEC OMVs, which also showed a significant decrease in the rate of uptake (Figure 6). This indicated that in contrast to MG1655, EHEC OMVs preferentially utilize clathrin-independent entry routes. Entry was also dynamin-dependent, suggesting OMVs containing O-antigen are taken up by caveolin-mediated endocytosis. OMVs from EHEC contribute to infection via delivery of diverse virulence factors, and this may be a further adaptation to pathogenesis for EHEC, as although the rate of internalization by caveolae is around 5 times slower than that of clathrin-dependent endocytosis, unlike clathrin-coated pits caveolae do not enter endocytic trafficking routes, and thus are not targeted to lysosomes for processing or degradation (Ritter et al, 1995; Bielaszewska et al, 2013). Instead, the contents of caveolae rapidly diffuse into the cytoplasm, consistent with our observation of a rapid increase in cytoplasmic dye cleaveage and thus, rapid decrease in FRET almost immediately after vesicle addition to cells. These findings underpin the notion that internalization via caveolin-mediated endocytosis allows more efficient delivery of cargo into host cells (Ritter et al, 1995). When clathrin-mediated endocytosis is inhibited, it becomes obligatory for OMVs to utilize the more efficient caveolin pathway, which would explain the increased total change in blue fluorescence observed for EHEC OMVs when cells are treated with papain or chlorpromazine. It may be that OMVs from MG1655 lacking O-antigen are less well adapted to using this route, and the presence of the O-antigen on OMVs from MG1655 L9 enhances their ability to use the alternative caveolin-mediated route of entry (Figure 7).

**Figure 7.**
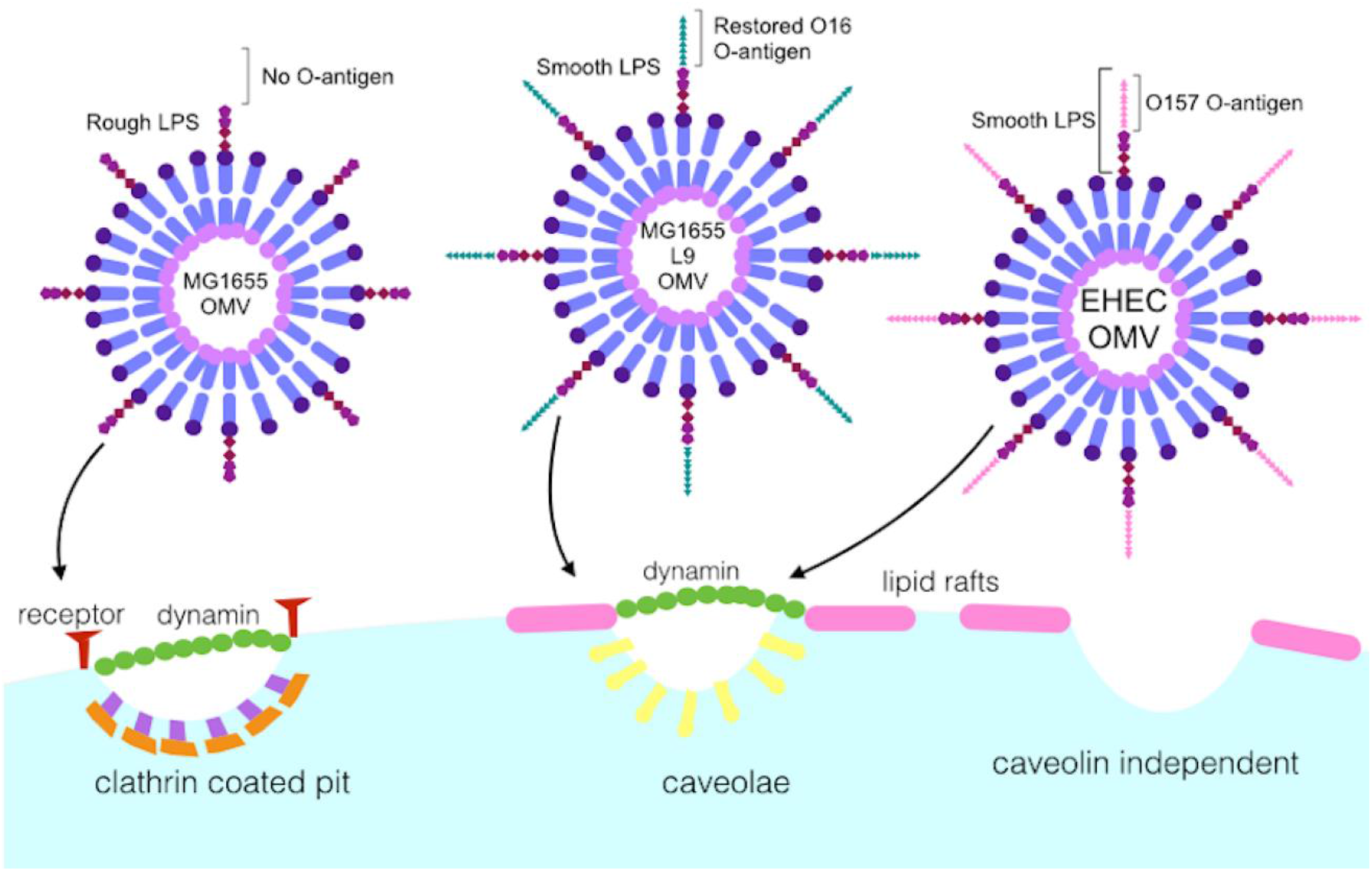
LPS composition determines major route and kinetics of OMV entry into host cells.

Previous work with *H. pylori* OMVs had indicated an inhibitory role of extracellular LPS on OMV uptake (Parker et al, 2010). In our experiments, we found that addition of purified LPS resulted in significant inhibition of uptake of EHEC OMVs, but had no effect on uptake of OMVs from MG1655 or L9. LPS is recognized by the TLR4 receptor on the host cell membrane, but activation of this receptor requires binding of LPS to the CD14 protein which is associated with lipid raft microdomains and caveolae (Shuto et al, 2005; Plociennikowska et al, 2015). EHEC OMVs were strongly dependent on lipid rafts for their entry, and were most affected by inhibition of this pathway with cholesterol sequestering agents (Figure 6). Inhibition of TLR4 using C34, a small molecule which competitively binds the TLR4-MD2 complex, had no significant effect on uptake of OMVs (Figure S4) (Neal et al, 2013). In addition, we showed that removal of surface receptors resulted in an increased rate of uptake for EHEC OMVs, suggesting they are not dependent on receptor-mediated endocytosis (Figure 6). We conclude that EHEC OMVs are competing with purified LPS for cholesterol-rich domains of the cell membrane required both for the formation of TLR4 signaling complexes and vesicular uptake, rather than directly competing for the TLR4 receptor.

Whilst it is well established that pathogenic species utilize OMVs during infection, the specific adaptations which allow OMVs to contribute to pathogenesis require further exploration. This work has developed a new approach to overcome current methodological limitations and provide consistent data for future studies and allow new insights into the interactions of OMVs with host cells during infection. Our data demonstrate the relevance of LPS composition, in particular the presence and chemical composition of O-antigen, in shaping route and entry kinetics of OMVs. Further work in this area may reveal targets for inhibition of these processes, and enable attenuation of infections by preventing the OMV-associated delivery of virulence factors.

## AUTHOR CONTRIBUTIONS

E.J.O, D.B., L.J.A. and A.M.K. designed experiments, D.B. and L.J.A contributed reagents, E.J.O. and E.B. performed experiments, E.J.O., E.B., S.J. and A.M.K. analyzed data, E.B. performed electron microscopy, and E.J.O., S.J. and A.M.K. contributed to writing the manuscript. All authors checked and approved the final version of the manuscript.

## ACKNOWLEDGMENTS

We thank Matthew DeLisa (Cornell Univ.) for providing ClyA-Bla and Bla-ClyA plasmid constructs and Peter Feng (FDA, Maryland) for providing the EHEC MA6 strain. We thank staff at the Birmingham Electron Microscopy Facility for technical help with sample preparation and transmission electron microscopy. We thank members of the Krachler lab for critical reading and comments on the manuscript. This work was supported by BBSRC grants BB/M021513/1 and BB/L007916/1 (to A.M.K.), and a BBSRC MIBTP studentship (to E.J.O).

## METHODS

### Bacterial strains and growth conditions

Enterohaemorrhagic *Escherichia coli* (EHEC) Sakai O157:H7, EHEC MA6 (provided by Peter Feng, FDA, Maryland) (Rump et al, 2010) *E. coli* MG1655 and *E. coli* MG1655 L9 with restored O-antigen (Browning et al, 2013) were transformed via electroporation with plasmid pBAD ClyA-Bla (provided by Matthew DeLisa, Cornell University), (Kim et al, 2008). The strains were grown in lysogeny broth (Miller, Sigma), with 50 μg/ml kanamycin, at 37 °C with shaking at 200 rpm.

### Isolation of outer membrane vesicles by ultracentrifugation

100 ml cultures were grown in LB at 37 °C, with agitation at 200 rpm. Once the OD_600_ reached 0.5-0.6, cells were induced with 0.2% L-arabinose and grown for a further 16 h. Cells were then pelleted at 6000xg, and the supernatants were removed and filtered with a 0.45um syringe filter. Aliquots of filtered supernatants were spread on LB agar and grown overnight at 37 °C to check that all viable cells had been removed by filtration. 25 ml of filtered supernatants were centrifuged in a Beckman XL90 ultracentrifuge using a 70Ti rotor at 100,000xg (30,000 rpm) for 2 h at 4 °C. After centrifugation, supernatants were removed, and the OMV pellets were resuspended in 1 ml colorless DMEM or sterile water (for TEM) and stored at −20 °C.

### Isolation of outer membrane vesicles by ExoSpin columns

Strains were grown in 5 ml LB containing 50 μg/ml kanamycin, and induced with 0.2% L-arabinose. Cultures were incubated for 18 h at 37 °C, with shaking at 200 rpm. Cells were pelleted at 6000xg for 5 min, and 1 ml supernatant was then transferred to new tubes and centrifuged at 17,200xg for 30 min at 4 °C. 100 μl of supernatant were then added to prepared exospin columns (Cell Guidance Systems) and centrifuged at 50xg for 1 min at 4 °C, and then OMVs were eluted by addition of 100μl PBS to the column and centrifuged at 50xg as before. The purified OMVs were then analyzed for size and concentration using nanoparticle tracking analysis.

### Nanoparticle tracking analysis

After purification, OMV samples were diluted 1x10^−6^ in filtered sterile PBS. Particle diameter and concentration were measured using the Nanosight LM10 particle tracking analysis, with a minimum of 100 tracks per sample, performed in triplicate. Camera shutter 1495 and gain of 450 were used, and size distribution scatter plots were created using GraphPad Prism. Size distribution was analysed using analysis of variance (ANOVA) with a Brown Forsythe test for equal variance.

### Visualization of outer membrane vesicles by Transmission Electron Microscopy

10μl of isolated outer membrane vesicles in sterile deionized distilled water were added to 400-mesh copper grids, and negatively stained with 4% uranyl acetate for 2 min. Samples were then observed using a Jeol 1200Ex transmission electron microscope (Birmingham Electron Microscopy Facility) with an acceleration of 75kV.

### Transmission Electron Microscopy of HeLa cells incubated with outer membrane vesicles

HeLa cells (P5) were seeded onto 13 mm coverslips at a concentration of 1x105 cells per ml in complete DMEM, 24 h prior to infection. The following day, OMVs were added to the cells for a final concentration of 10 μg/ml protein. The infection proceeded for 0, 30 or 60 min before coverslips were placed in 2.5% glutaraldehyde fixative, and stored at 4 °C before processing and embedding (Birmingham Electron Microscopy Facility). The samples were observed using the Jeol 1200 transmission electron microscope, with acceleration at 70kV.

### Western blotting of EHEC cellular fractions

12 μl of samples normalized for their protein content from EHEC ClyA-Bla and Bla-ClyA whole cell lysate, supernatant and OMV fractions were added to 3μl 5X SDS loading dye and boiled for 10 min. Samples were loaded onto a 15 well BioRad pre-cast stain-free SDS-PAGE gel and run at 120V, 200mA for 45 min. The gel was then transferred onto a PVDF membrane in transfer buffer containing 20% methanol for 80 minutes at 100V. After transfer, the membrane was blocked at room temperature in TBS 0.1% Tween-20 and 5% skim milk for 1h with agitation. The membrane was washed 3 times with TBS 0.1% Tween-20 (5 min per wash). After blocking, the membrane was incubated with a 1:2000 dilution of mouse anti-Bla primary antibody in TBS 0.1% Tween-20 and 5% skim milk overnight at 4 °C with agitation. The following day, the membrane was washed 3 times as before, and incubated with a 1:5000 dilution of sheep anti-mouse secondary antibody in TBS 0.1% Tween-20, 5% skim milk for 1h at room temperature with agitation. The membrane was washed again 3 times, and 2 ml BioRad ECL reagents were added to the membrane and incubated for 5 min, before visualization with a BioRad ChemiDoc imager.

### Protein Quantitation

To quantify levels of protein in cell fractions, the ThermoFisher CBQCA Protein Quantitation kit was used. 10 μl of protein was added to 125 μl 0.1M sodium borate, 5 μl 20 mM KCN, and 10 μl 5 mM ATTO-TAG in a 96-well plate, and incubated in the dark for 1-2 h. The fluorescence in samples was measured with an excitation at 465 nm and an emission at 550 nm in a FluoStar OMEGA plate reader. The level of fluorescence in samples was compared to known quantities of bovine serum albumin, and this was used to determine and normalize the protein concentration in cellular fractions.

### Nitrocefin assay to determine β-lactamase activity

50 μl of samples were added in triplicate to a 96-well plate. Nitrocefin was diluted to 0.5 mg/ml in PBS and 50 μl was added to each sample. The absorbance at 486 nm was measured in the FluoStar Omega plate reader for 2 h, and the change in absorbance over time was used to determine the specific activity in samples, using the protein concentration determined by the CBQCA kit.

### Papain and detergent treatment of OMVs

Triton X-100 and SDS were added at a concentration of 1% to 20 μl OMVs for 45 min at 37 °C. 5ug/ml papain was then added for 30 or 60 min at 37 °C. The papain reaction was inactivated using 1 mM PMSF at room temperature for 30 min. 5 μl SDS-PAGE loading dye was added to the samples, which were then boiled for 10 min. Samples were run on a 15-well pre-cast stain free gel for 45 min at 120V, and then subjected to Western blotting with anti-β-lactamase primary antibody (Pierce) as described above.

### LPS staining of OMVs

OMV concentrations were normalised, and 50ul of samples were treated with 5μg/ml proteinase K (Sigma) for 1h at 60 °C to generate LPS only samples. 15μl of treated OMVs were then run on a pre-cast BioRad 12% acrylamide gel at 120V for 1h. The gel was then fixed and stained using the Pierce Silver Stain kit, or the Thermo ProQ Emerald staining kit according to the manufacturers instructions.

### Cellulase treatment of OMVs

1% cellulase (Sigma) was added to EHEC ClyA-Bla OMVs for 2h at 37 °C, and OMVs were subsequently re-purified by ultracentrifugation.

### Plate reader FRET experiments

HeLa cells (passage 1-7) were seeded in triplicate in a black-walled, clear bottom 96-well plate at a concentration of 1x10^5^ cells per ml in Dulbecco’s modified Eagle medium (DMEM) supplemented with 1% L-glutamine, 1% Penicillin/Streptomycin and 10% heat inactivated fetal bovine serum. The plate was incubated at 37 °C, 5% CO_2_ for 24 h prior to experiments. The following day, cells were loaded with 20 μl 6X CCF2-AM with 100 μl colourless unsupplemented DMEM (cDMEM) and incubated at room temperature for 1 h in the dark to allow dye loading. The dye was removed by washing 2x in PBS and 1x in cDMEM. Cells were treated with 5 mM methyl-ß-cyclodextrin or 1 μg/ml filipin to inhibit cholesterol mediated endocytosis, 80 uM Dynasore for dynamin inhibition, or 20 uM blebbistatin for macropinocytosis inhibition for 1h at 37 °C. Cells were treated with 1 μg/ml chlorpromazine for 1h at 37 °C to inhibit formation of clathrin-coated pits, or with 5 μg/ml papain for 15 min at 37 °C to remove surface proteins, before inactivation of papain with 5 mM PMSF for 20 min. Rough LPS (from *E. coli* EH100 Ra mutant strain, Sigma) or smooth LPS (from *E. coli* O55:B5, Sigma) was added at a concentration of 1μg/ml, and TLR4 inhibitor C34 was added at a concentration of 10μM for 30 min at 37 °C prior to co-incubation of OMVs and cells.

Reporter OMVs were diluted in cDMEM and added to the cells for a final concentration of 10 μg/ml, or 1x10^8^ vesicles, corresponding to an MOI of 1000. The plate was immediately placed in the PheraStar plate reader, with excitation at 405 nm and simultaneous dual emission at 530 nm and 460 nm. The wells were scanned (bottom optic) with orbital averaging for a total of 150 cycles, equating to a measurement every 90 seconds for 3 hours. The ratio of blue to green fluorescence intensity detected in the cells at each cycle was calculated using GraphPad Prism, and ratios for uninfected, dye-loaded cells were used as the baseline value for each cycle. All traces were normalized to 0 for their first ratio value. All experiments were performed with a minimum of three technical replicates and three independent repeats.

### Efficiency of uptake and statistical analysis

Efficiency of uptake was calculated as the total change in blue:green fluorescence intensity ratio after 3h. Student’s t-test or ANOVA was used to determine statistical significance of total change between samples. A p value of <0.05 was considered statistically significant.

### Rate estimation and statistical analysis

To estimate the gradients of the data, polynomials were fitted to each data set using the cubic spline function *csaps* in Matlab. Numerical estimates of the gradients of the resulting polynomials were determined using the *gradient* function. To ensure that the gradient estimates were as smooth as possible whilst also retaining the overall shape and trend of the data, a small smoothing parameter was used.

ANOVA was used to analyse data sets, with a Brown Forsythe test to determine equal variance using GraphPad Prism software. A p-value of <0.05 was considered statistically significant.

### Confocal Microscopy

HeLa cells (P3-7) were seeded on 13mm coverslips in a 12-well plate at a concentration of 1x105 cells per ml in complete DMEM, 24 h prior to experiments. The following day, cells were washed and loaded with 100 μl 6X CCF2-AM dye with 500 μl colourless unsupplemented DMEM, and incubated in the dye solution for 1 h at room temperature in the dark. Cells were then incubated with ClyA-Bla reporter OMVs for 0-4 h. The cells were washed with PBS and then fixed with 0.5 ml 4% PFA for 15 min. The next day, coverslips were mounted onto slides with a drop of Gold Anti-Fade mounting solution and then imaged using the Nikon A1R confocal microscope (Birmingham Advanced Light Microscopy Facility), and fluorescence was observed from excitation at 409 nm and dual emissions at 450 nm and 520 nm. Z stacks were produced from 20 slices, with gain, exposure and laser intensity kept the same for all slides, and images were taken for 3 representative fields of view per slide. The Z stacks were converted to maximum intensity projection images.

## Supplemental Information

**Figure S1.**
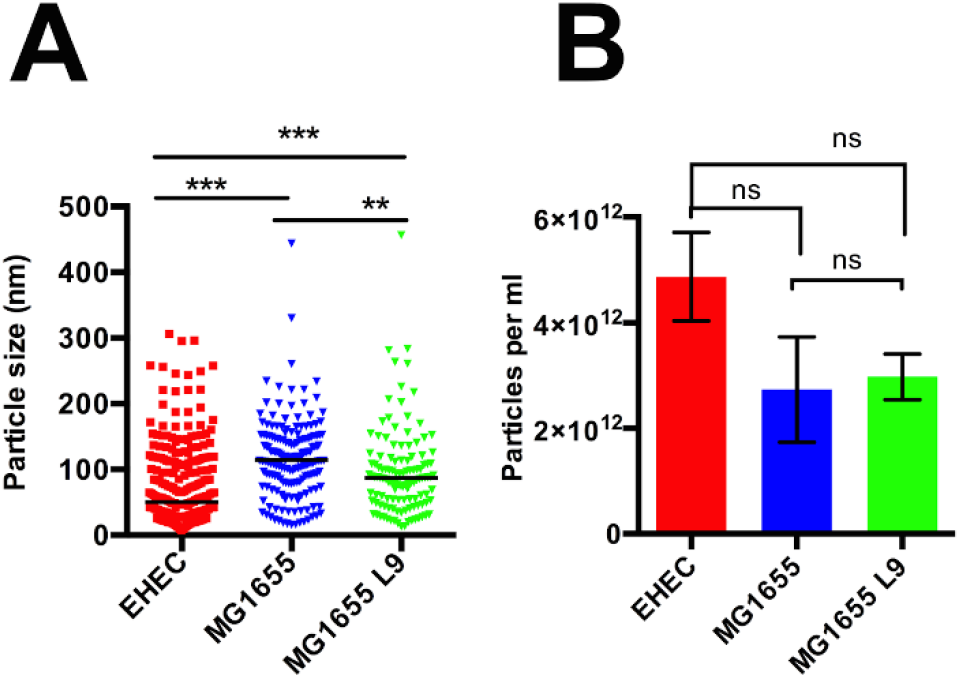
Size distribution and concentration of OMVs. (A) OMVs isolated from EHEC ClyA-Bla (red), MG1655 ClyA-Bla (blue) and MG1655 L9 ClyA-Bla (green) were diluted 1x10^−6^ and analyzed using nanoparticle tracking analysis to determine their size distribution and concentration. Data represents individual tracks, and means ± stdev from at least 200 tracks per sample. Significance was determined using analysis of variance (ANOVA), with a Brown Forsythe test to determine equal variance. (***) indicates p value of < 0.001, (**) p <0.01. (B) Particle concentrations of OMV preparations isolated from EHEC ClyA-Bla (red), MG1655 ClyA-Bla (blue) and MG1655 L9 ClyA-Bla (green) were measured using nanoparticle tracking analysis, then used to dilute OMVs to generate equal MOIs of 1000 (10^8^ OMVs/ml) for infection experiments. Results are means ± stdev from at least 200 tracks per sample. Significance was determined using analysis of variance (ANOVA), with a Brown Forsythe test to determine equal variance. NS indicates no significant differences between samples (p value > 0.05).

**Figure S2.**
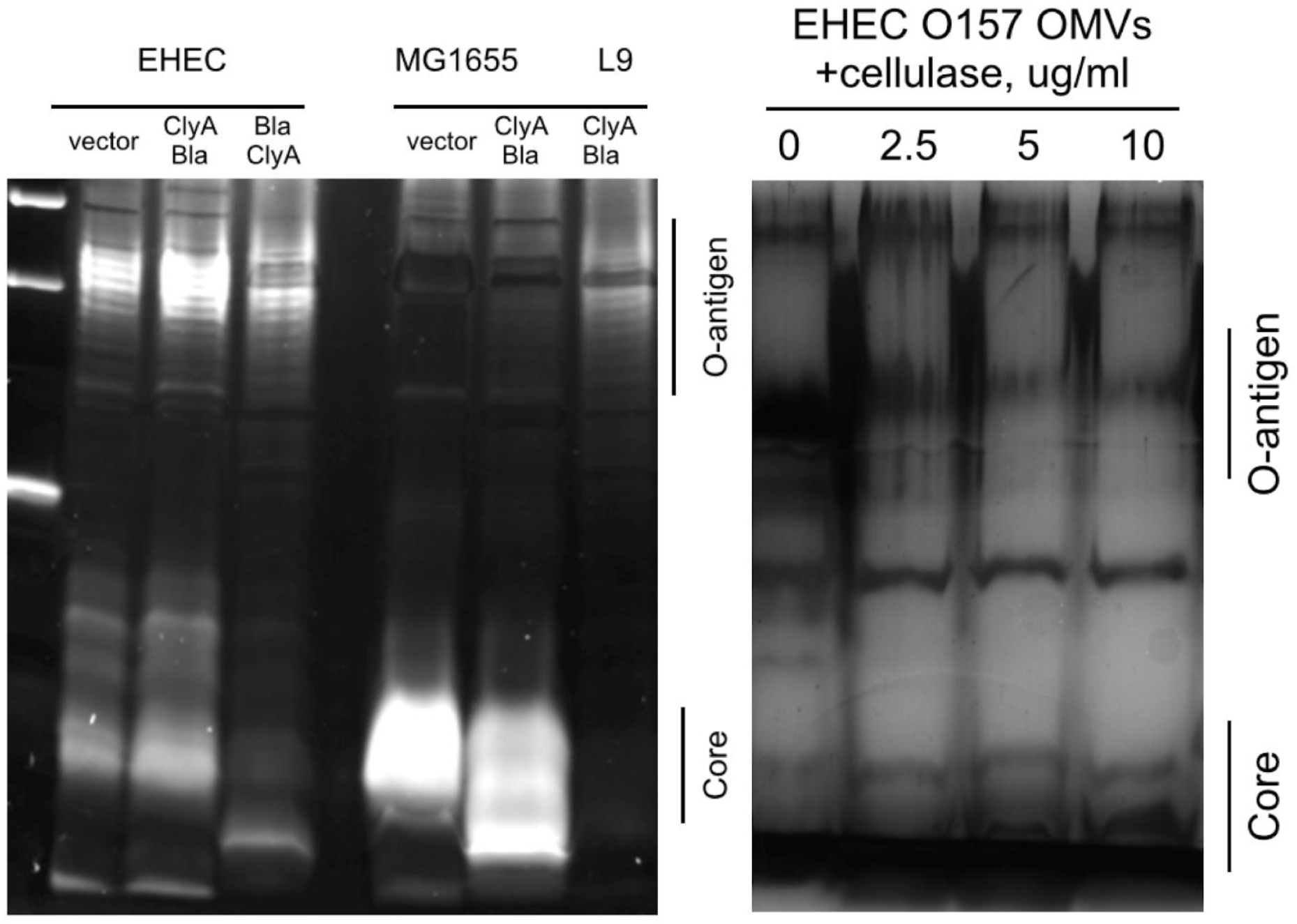
Characterization of lipopolysaccharide from EHEC wild type, MA6, MG1655 and MG1655 L9 strains. OMVs were treated with proteinase K to generate LPS-only samples. EHEC OMVs were also treated with varying concentrations of cellulase to enzymatically remove O-antigen. Isolated LPS was run on pre-cast SDS-PAGE gels and silver stained. High molecular weight bands indicate presence of O-antigen.

**Figure S3.**
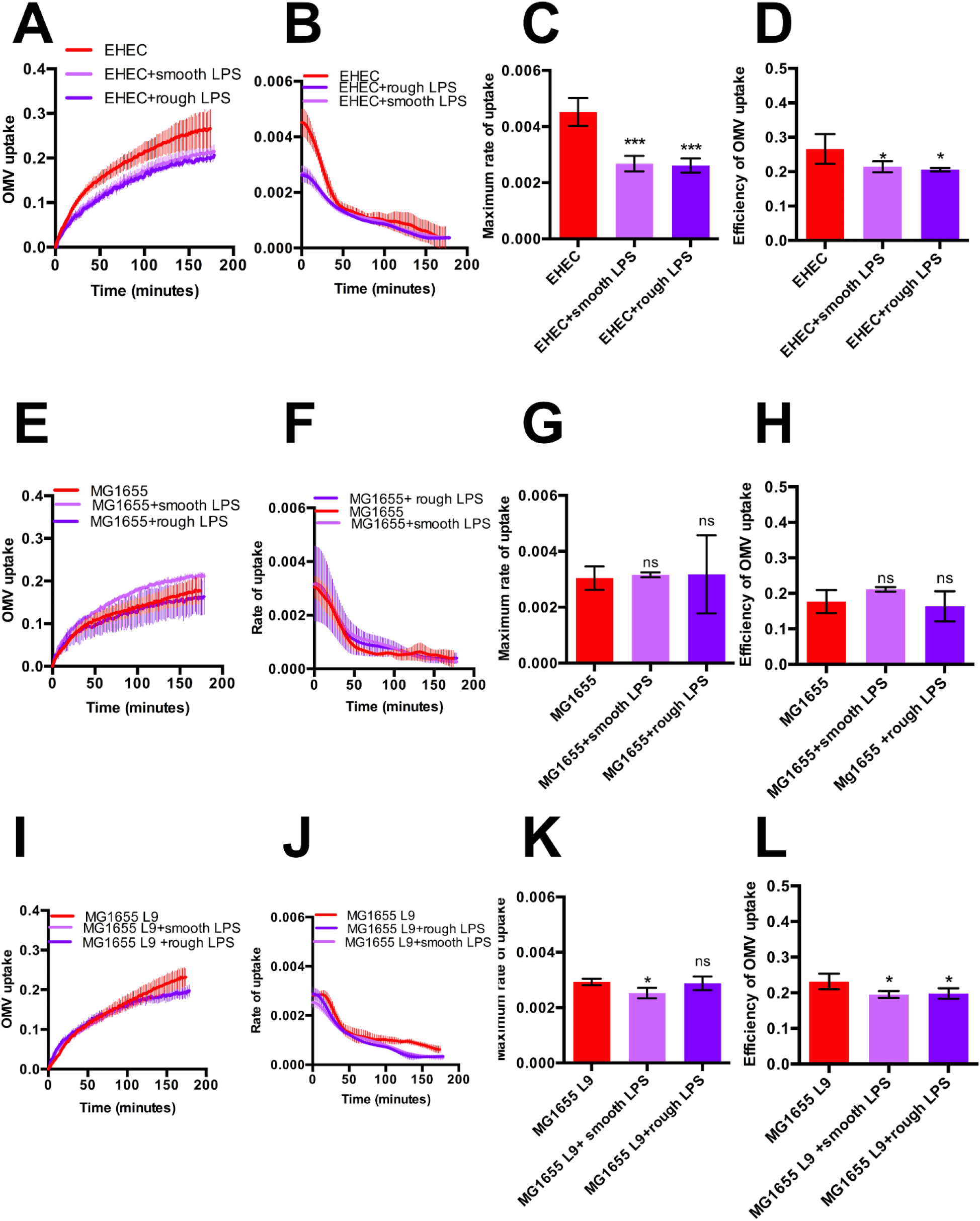
Supplementation with exogenous LPS inhibits entry of EHEC OMVs. (A) Hela cells were either left untreated (red), or pre-treated with 1ug/ml smooth LPS (lilac) or 1ug/ml rough LPS (violet) and exposed to ClyA-Bla OMVs isolated from EHEC at an MOI of 1000 for 3 hours. The FRET signal (ratio of blue:green fluorescence) over time was plotted as mean ± stdev (n=3). (B) Polynomials were fitted to each data set from (A) using the cubic spline function csaps in Matlab. Numerical estimates of the gradients of the resulting polynomials were determined using the gradient function and plotted as means ± stdev (n=3). (C) Maximum rates were determined from data in (B) to visualize speed of OMV uptake into untreated Hela cells (red), or cells pre-treated with 1ug/ml smooth LPS (lilac) or 1ug/ml rough LPS (violet). Data shown are means ± stdev (n=3) and significance was determined using ANOVA (****) indicates p≤0.0001, ns-not significant (p≥0.05). (D) Total FRET changes after 3 hrs were determined from data in (A) and plotted to visualize overall efficiency of uptake into untreated Hela cells (red), or cells pre-treated with 1ug/ml smooth LPS (lilac) or 1ug/ml rough LPS (violet). Data shown are means ± stdev (n=3) and significance was determined by ANOVA (***) indicates p≤0.001, (**)p≤0.01. (E) Hela cells were either left untreated (red), or pre-treated with 1ug/ml smooth LPS (lilac) or 1ug/ml rough LPS (violet) and exposed to ClyA-Bla OMVs isolated from MG1655 at an MOI of for 3 hours. The FRET signal (ratio of blue:green fluorescence) over time was plotted as mean ± stdev (n=3). (F) Polynomials were fitted to each data set from (A) using the cubic spline function csaps in Matlab. Numerical estimates of the gradients of the resulting polynomials were determined using the gradient function and plotted as means ± stdev (n=3). (G) Maximum rates were determined from data in (B) to visualize speed of OMV uptake into untreated Hela cells (red), or cells pre-treated with 1ug/ml smooth LPS (lilac) or 1ug/ml rough LPS (violet). Data shown are means ± stdev (n=3) and significance was determined using ANOVA (****) indicates p≤0.0001, ns-not significant (p≥0.05). (H) Total FRET changes after 3 hrs were determined from data in (A) and plotted to visualize overall efficiency of uptake into untreated Hela cells (red), or cells pre-treated with 1ug/ml smooth LPS (lilac) or 1ug/ml rough LPS (violet). Data shown are means ± stdev (n=3) and significance was determined by ANOVA (***) indicates p≤0.001, (**)p≤0.01. (I) Hela cells were either left untreated (red), or pre-treated with 1ug/ml smooth LPS (lilac) or 1ug/ml rough LPS (violet) and exposed to ClyA-Bla OMVs isolated from MG1655 L9 at an MOI of 1000 for 3 hours. The FRET signal (ratio of blue:green fluorescence) over time was plotted as mean ± stdev (n=3). (J) Polynomials were fitted to each data set from (A) using the cubic spline function csaps in Matlab. Numerical estimates of the gradients of the resulting polynomials were determined using the gradient function and plotted as means ± stdev (n=3). (K) Maximum rates were determined from data in (B) to visualize speed of OMV uptake into untreated Hela cells (red), or cells pre-treated with 1ug/ml smooth LPS (lilac) or 1ug/ml rough LPS (violet). Data shown are means ± stdev (n=3) and significance was determined using ANOVA (****) indicates p≤0.0001, ns-not significant (p≥0.05). (L) Total FRET changes after 3 hrs were determined from data in (A) and plotted to visualize overall efficiency of uptake into untreated Hela cells (red), or cells pre-treated with 1ug/ml smooth LPS (lilac) or 1ug/ml rough LPS (violet). Data shown are means ± stdev (n=3) and significance was determined by ANOVA (***) indicates p≤0.001, (**)p≤0.01.

**Figure S4.**
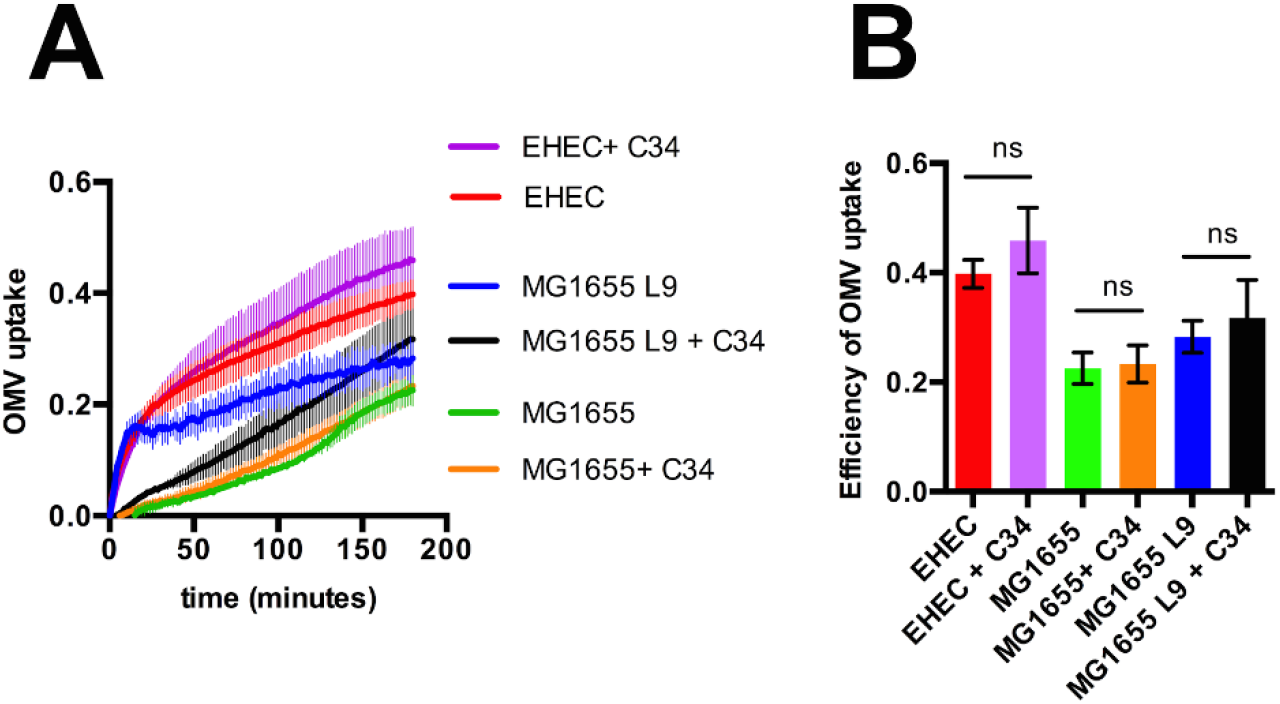
Inhibition of TLR4 has no effect on OMV uptake. (A). Hela cells were either left untreated (red, green and blue for O157, MG1655 and MG1655 L9 respectively, or pre-treated with 10uM TLR4 inhibitor C34 (lilac, orange and black for O157, MG1655 and MG1655 L9 respectively) exposed to ClyA-Bla OMVs isolated from O157, MG1655 and MG1655 L9 at an MOI of 1000 for 3 hours. The FRET signal (ratio of blue:green fluorescence)1046over time was plotted as mean ± stdev (n=3). (B). Total FRET changes after 3 hrs were determined from data in (A) and plotted to visualize overall efficiency of uptake into untreated Hela cells (red), or cells pre-treated with C34. Data shown are means ± stdev (n=3) and significance was determined by ANOVA. (ns) indicates p value > 0.05.

